# Molecular evolution of classic Hodgkin lymphoma revealed through whole genome sequencing of Hodgkin and Reed Sternberg cells

**DOI:** 10.1101/2021.11.05.467496

**Authors:** Francesco Maura, Bachisio Ziccheddu, Jenny Z. Xiang, Bhavneet Bhinder, Federico Abascal, Kylee H. Maclachlan, Kenneth Wha Eng, Manik Uppal, Feng He, Wei Zhang, Qi Gao, Venkata Yellapantula, Sunita Park, Matthew Oberley, Elizabeth Ruckdeschel, Megan S. Lim, Gerald Wertheim, Matthew Barth, Terzah M. Horton, Christopher Forlenza, Yanming Zhang, Ola Landgren, Craig H Moskowitz, Ethel Cesarman, Marcin Imielinski, Olivier Elemento, Mikhail Roshal, Lisa Giulino-Roth

## Abstract

The rarity of malignant Hodgkin and Reed Sternberg (HRS) cells within a classic Hodgkin lymphoma (cHL) biopsy limits the ability to study the genomics of cHL. To circumvent this, our group has previously optimized fluorescence-activated cell sorting to purify HRS cells. Here we leveraged this method to report the first whole genome sequencing landscape of HRS cells and reconstruct the chronology and likely etiology of pathogenic events prior to the clinical diagnosis of cHL. We identified alterations in driver genes not previously described in cHL, a high activity of the APOBEC mutational signature, and the presence complex structural variants including chromothripsis. We found that the high ploidy observed in cHL is often acquired through multiple, independent large chromosomal gain events including whole genome duplication. The first of these likely occurs several years prior to the diagnosis of cHL, and the last gains typically occur very close to the time of diagnosis. Evolutionary timing analyses revealed that driver mutations in *B2M, BCL7A, GNA13*, and *PTPN1*, and the onset of AID driven mutagenesis usually preceded large chromosomal gains. The study provides the first temporal reconstruction of cHL pathogenesis and suggests a relatively long time course between the first pathogenic event and the clinical diagnosis.

## Article

Classic Hodgkin lymphoma (cHL) is characterized by a unique pathological composition where a small fraction of Hodgkin and Reed Sternberg (HRS) tumor cells (∼1%) are surrounded by an extensive and complex immune and stromal infiltrate.^1^ The paucity of HRS cells in tumor tissue has precluded the genomic investigation of cHL using standard platforms. Our group has optimized fluorescence-activated cell sorting (FACS) to isolate HRS cells and intratumor B- and T-cells and perform whole exome sequencing (WES).^2^ Data on HRS exomes from our group and others have revealed mutations in critical driver genes involved in NF-κB and JAK/STAT signaling pathways as well as genes involved in immune escape.^2-5^ These investigations, however, were limited to captured coding sequences, and therefore not able to comprehensively decipher the genomic complexity of cHL. Furthermore, the chronologic order in which somatic alterations are acquired is largely unknown in cHL, limiting our understanding of the earliest drivers/initiating events.

Whole genome sequencing (WGS) has the potential to fully characterize the somatic genomic landscape including: i) the catalogue of coding and non-coding mutations, ii) large and focal copy number alterations (CNA), iii) structural variants (SV) including complex events, and iv) mutational processes involved in cancer pathogenesis.^6^ Here we performed WGS on FACS-isolated HRS cells and matched normal tissue from 25 patients with cHL and WES from an additional 36 cases. We combined CNA, single nucleotide variant (SNV) and SV analyses to decipher the landscape and chronological order of key driver events in cHL. Altogether these analyses reveal previously unknown insights into the pathogenesis of cHL and pave the way for the design of novel therapeutic strategies targeting early driver events in a disease that frequently affects children and young adults.

## Results

### Single nucleotide variants in classical Hodgkin lymphoma driver genes

To evaluate the genome of HRS cells, HRS and intra-tumoral T-cells were isolated from cHL biopsies using FACS as previously described.^7^ Intra-tumoral T-cells from each case were used as the germline control. We interrogated WGS from 25 cases of cHL including 10 pediatric cases (age<18), 9 cases in adolescents and young adults (AYA, age 18-40), and 6 in older adults (age>40) (**Supplemental Table 1**). Twenty-two (88%) of the cases were obtained at the time of diagnosis and 3 (12%) were obtained at the time of relapse. The median DNA input from HRS cells was 13.6ng (range 4.2 - 226ng). Given the ultra-low input, sequencing data were generated by amplifying the DNA (median 10 amplification cycles, range 7-15). To ensure a robust quality control, we explored possible amplification-based mutational artifacts across the genome, observing an enrichment of single base substitution (SBS) within distinct trinucleotide context reflecting palindromic artifacts (**Supplemental Figure 1** and **Supplemental Table 2**).^8,9^ After having removed these amplification-induced palindromic sequencing artifacts, we observed a median mutational burden of 5006 per genome (range 1763-18436), which is comparable to other aggressive lymphomas (**Figure 1a**).^10^ We found that pediatric and AYA patients age ≤40y had a significantly higher mutational burden than older adults over age 40 (median 5279 vs. 2945, p=0.009, **Figure 1b**). This effect was not related to the sequencing coverage (p=0.1).

**Figure 1.**
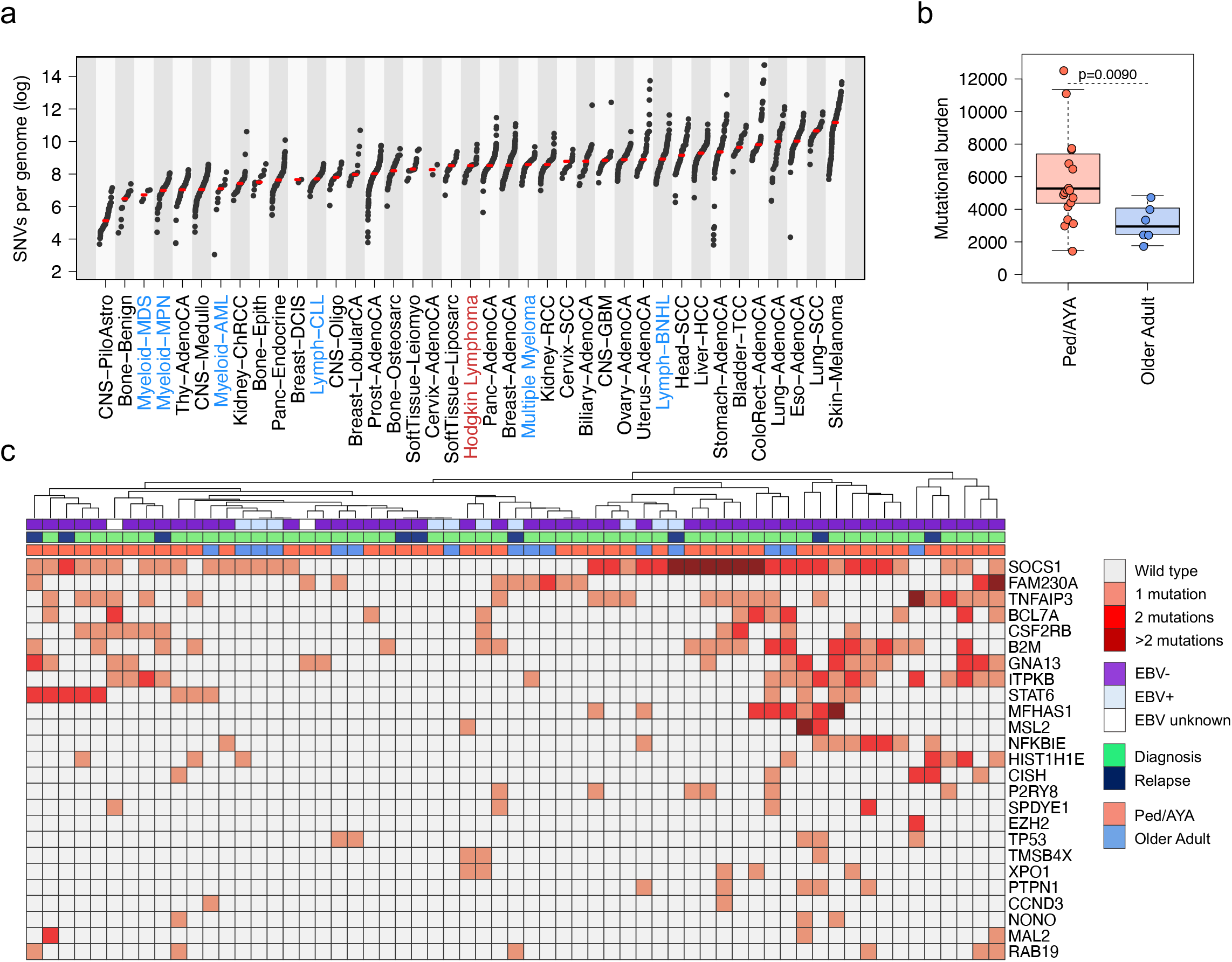
cHL mutational landscape. **a)** WGS mutational burden comparison between cHL and other cancers included in the PCAWG (n= 2780) and in MM (n=71) WGS studies. cHL is highlighted in red, other hematological cancers in blue. The median of each tumor type is annotated with a red line. **b)** Whole genome mutational burden comparison between Ped/AYA and older adults with cHL. p-value was calculated using Wilcoxon rank-sum test. Boxplots show the median and interquartile range. c) Heatmap summarizing the 25 mutated driver genes/hot spots across 61 cHL patient with available either WGS or WES data.

To increase the sample size for discovery analysis of driver mutations, we performed WES on HRS cells from additional 36 cHL cases, 10 of which have been previously reported^2^ (**Supplemental Table 3**). After having identified and removed the same amplification-induced sequencing palindromic artifact, we combined the two cohorts [WGS (n=25) WES (n=36)] to perform a driver mutation discovery analysis. To identify genes that are hit by nonsynonymous mutations more frequently than what would be expected by chance (i.e. genes under positive selection), we ran *dndsv*^11^ both on all genes and, to increase statistical power, on a restricted set of cancer genes built by combining the COSMIC census catalogue with genes previously reported as recurrently mutated in cHL (n=15).^2-5^ Overall, *dndscv* detected 23 genes under positive selection (q<0.1) and five hotspots (*CCND3, TP53, B2M, EZH2*, and *XPO1*) (**Figure 1c, Supplemental Table 4-5**). Ninety-five percent of cHL cases had at least one of these driver genes mutated, with an average of 3.38 (range 2.45-3.98) drivers per sample. Pediatric/AYA and older adult cases did not differ in the estimated number of mutations in driver genes calculated based on the global ratio of nonsynonymous to synonymous mutations.

The most common driver alterations were in *SOCS1* (62% of cases), *TNFAIP3* (36%) and *B2M* (32%). We observed alterations in 11 driver genes not previously described in cHL (**Supplemental Table 4**),^2-5,12^ including *BCL7A* and *CISH. BCL7A* encodes a subunit of the SWI/SNF complex and has been described as a tumor suppressor in B-cell non-Hodgkin lymphomas.^13,14^ Cytokine inducible SH2 containing protein (*CISH*) is a member of the SOCS family which regulates cytokine receptor signaling through the JAK/STAT pathway.^15^

*B2M, BCL7A, GNA13, ITPKB*, and *SOCS1* tended to have more than one nonsynonymous mutation in the same patient (**Figure 1c**). Using the WGS mutational distribution on the entire footprint of these hypermutated genes, we observed that *BCL7A*, and *ITPKB* showed a higher non-coding:coding mutation ratio compared to other driver genes (FDR<0.1 in Fisher’s exact test). Most of these mutations were compatible with the AID mutational signature (AID/SBS84 COSMIC signature; **Supplemental Figure 2**).^10,16-18^ The *SOCS1* mutational profile was also driven by AID/SBS84 mutational activity however the non-coding:coding mutation ratio was not significantly higher compared to other drivers. The presence of localized AID hypermutation activity, likely related to somatic hypermutation (SHM), provides a mechanistic explanation of why these genes tend to be mutated multiple times in the same patient. In contrast, *B2M* and *GNA13* were recurrently mutated more than once without enrichment for either intronic or SHM/AID mutations (**Supplemental Figure 2**).

### Copy number alterations (CNA)

Combining WGS and WES data, we investigated the somatic CNA landscape in cHL. Consistent with the frequent multinucleated nature of HRS cells, we observed high ploidy (median 2.95, range 1.66-5.33). Despite the well characterized bi- and multi-nucleated phenotype of Reed Sternberg cells, not all patients had whole genome duplication (WGD, 54%). This was validated by FISH in six patients with available material (**Supplemental Figure 3**). This suggests that multinucleation is not universally associated with WGD in cHL.^19-21^

Running GISTIC2.0,^22^ which identifies genes and chromosomal segments recurrently targeted by somatic CNA, we identified 19 recurrent CNA peaks: 5 gains and 14 losses (**Figure 2a**, and **Supplemental Table 6-7**). The majority of these CNAs were caused by either large (>10 Mb) or whole chromosome/arm events (95.6%; **Supplemental Figure 4-5**). Matching GISTIC CNA peaks with genes that are either positively selected in cHL and/or reported in the COSMIC census, we identified 4 genes recurrently involved in amplifications and 17 by deletions. For example, in *TNFAIP3*, the most frequently deleted gene (44% of all patients), 77% of deletions were either large (>10 Mb) or whole arm/chromosome loss. The most frequently amplified loci were 9p24.1 (*PDCD1LG2*/*JAK2*/*CD274*), and 2p16.1 (*REL/XPO1*), (67% and 85% of patients respectively, **Supplemental Figure 4**). These amplifications were similarly mostly caused by large and whole chromosome/arm duplications (80% and 85% respectively). Overall, our data suggests that known cHL driver genes are recurrently involved in large CNA, and, in a smaller proportion, by focal events. We also observed a high prevalence of high-level gains (>6 copies) in 2p16.1 – *REL/XPO1* [median 11 copies (range 7-31); n=10, 16%] and 9p24.1 – *PDCD1LG2*/*JAK2*/*CD274* [median 8 copies, range 6.3-13); n=11, 18%] (**Figure 2b**).

**Figure 2.**
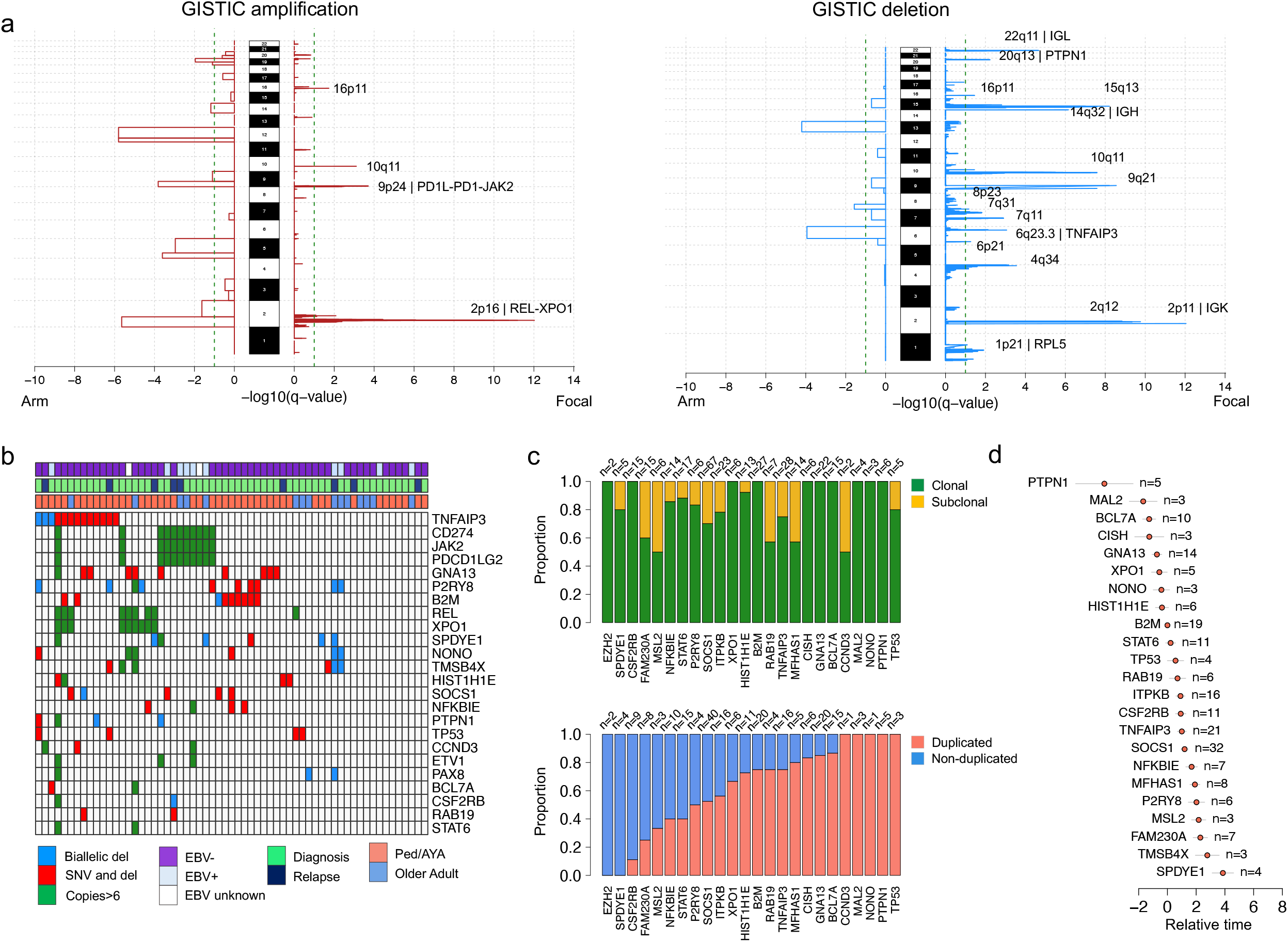
Copy number alterations (CNA) in cHL. **a)** Significant CNA GISTIC2.0 peaks and involved genes. **b)** Heatmap summarizing bi-allelic events and high level gains (>6 copies) involving driver genes extracted by GISTIC (n=21) and by *dndscv* (n=27). **c)** Clonal/subclonal and duplicated/non-duplicated distribution of mutations in driver genes. **d)** Chronological order of mutations in 25 cHL driver genes. Two out of 25 driver mutations are not reported because they had no relative chronological association with any other mutations in driver genes.

Next, we combined CNA data and nonsynonymous mutations to investigate biallelic events involving driver genes extracted by GISTIC2.0 (n=21) and by *dndscv* (n=25). *TNFAIP3* (n=15; 24%), *B2M* (n=10; 16%), *GNA13* (n=9; 15%) were the most common driver genes with bi-allelic inactivation. This was mostly driven by deletion on one allele and a nonsynonymous mutation on the other (**Figure 2b**).

### Timing of cHL driver alterations

Next, we investigated the relative timing of driver mutation acquisition with respect to chromosomal gains. For this analysis we included chromosomal gains of any size and level. By leveraging the high cHL ploidy and high prevalence of chromosomal gains, we performed a comprehensive investigation of the relative timing of driver mutation acquisition with respect to the chromosomal gain in 61 patients (**Figure 2c**).^23-26^ Clonal mutations within chromosomal gains can have one of two purity-corrected variant allelic frequencies (VAFs): i) ≤33% reflecting mutations acquired either on one of the two duplicated alleles after the gain or on the minor allele non-duplicated allele; ii) ≥66% reflecting mutations acquired on the duplicated allele before the duplication. A large fraction of mutations in driver genes (140/227; 61%) showed a duplicated VAF (≥66%), suggesting that they were acquired prior to chromosomal gains (**Figure 2c**). Loss of function mutations involved by copy neutral loss of heterozygosity (LOH) showed the same pattern, with the mutation often proceeding the chromosomal duplication. Next, we estimated the chronological order of mutations in 25 cHL driver genes/hot spots (**Figure 2d**). Temporal estimates were generated by the Bradley Terry model based on the integration between the CCF and duplication status of all mutations involving the 25 driver genes. Mutations in *PTPN1, GNA13, XPO1, HIST1H1E* and *B2M* emerged as early drivers occurring prior to other mutations and prior to chromosomal gain events.

### Mutational signatures

To investigate which mutational processes are involved in cHL, we used both *sigProfiler* and the *hierarchical Dirichlet process* (hdp) identifying 5 main mutational signatures [or single base substitution (SBS) signatures], and subsequently *mmsig* to accurately estimate their activity and contribution (**Figure 3a-b**).^10,17,26-29^ SBS1 and SBS5, the so-called clock-like mutational processes, were detected in all patients (median of 4250 mutations, range 735-11159).^30^ Seventy-two percent of patients had evidence of APOBEC mutational activity (SBS2 and SBS13), (**Figure 3b-c**). This high prevalence of APOBEC is similar to what has been reported based on HL WES data^5^ and confirms the strong APOBEC pathogenetic role in cHL. Four patients (16%) showed a particularly high APOBEC contribution mostly driven by a major APOBEC3A activity compared to APOBEC3B (hyper APOBEC), in line with what has been reported in other cancers.^26,31^ The other 14 (84%) were characterized an APOBEC3A:3B ratio ∼1 (“canonical” APOBEC). Looking at the APOBEC contribution before and after large chromosomal gains, we observed an enrichment among the non-duplicated mutations compared to the duplicated reflecting a later role in cancer pathogenesis (i.e., after the gain; p=0.002 using paired Wilcoxon test; **Figure 3d**).^32^ In contrast to other cancers like multiple myeloma,^23,26^ APOBEC activity was also detectable before the chromosomal gains, suggesting that large chromosomal duplications often occur in a clone in which APOBEC is already active.

**Figure 3.**
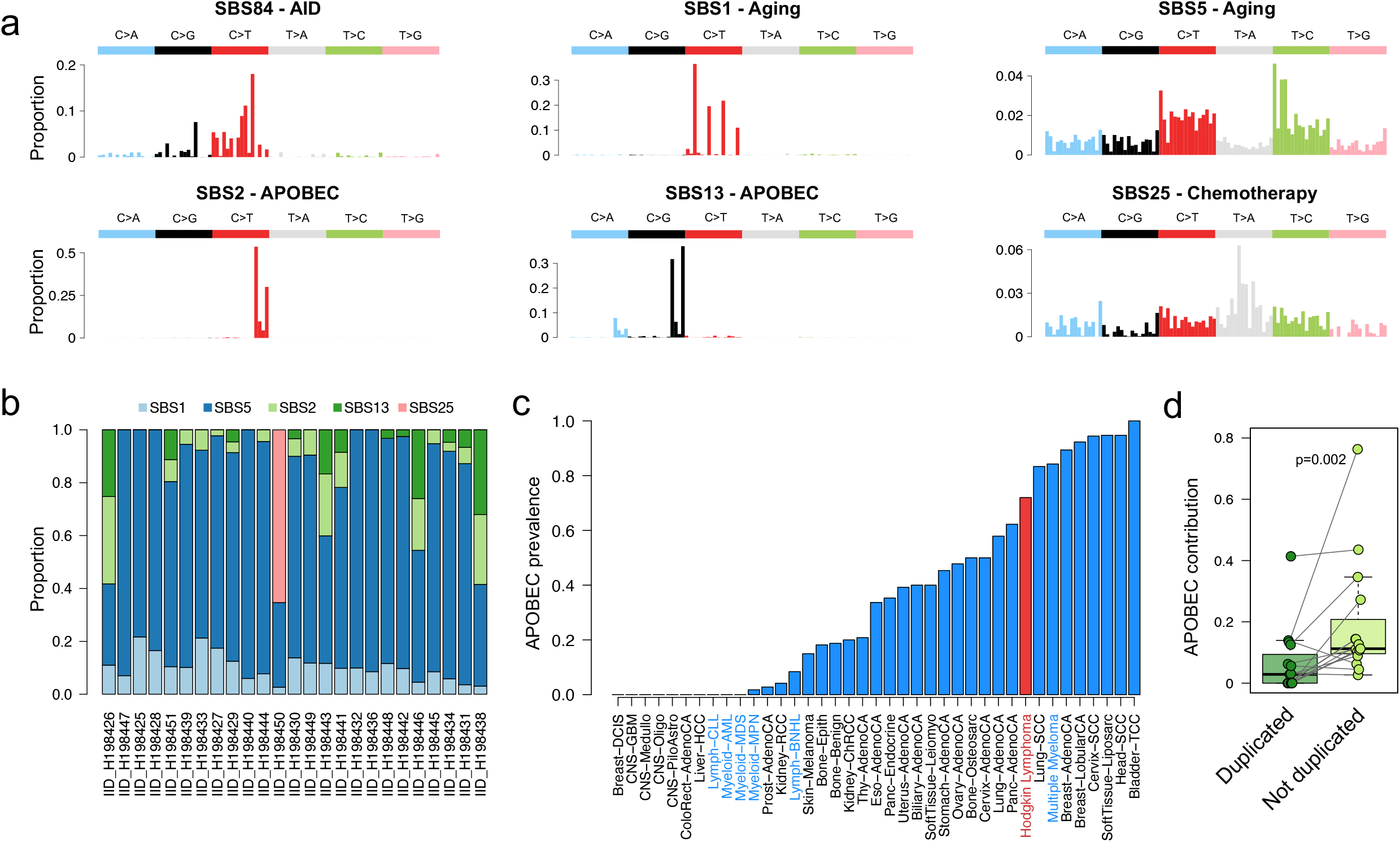
Mutational signatures in cHL. **a)** COSMICv.2 96-mutational profile of the 6 mutational signatures extracted and identified in cHL. **b)** Contribution of mutational signatures across 25 cHL cases with available WGS data. **c)** Proportion of patients with APOBEC activity across cHL and other cancers included in the PCAWG and in MM WGS studies. cHL is highlighted in red, other hematological cancers in blue. **d)** APOBEC relative contribution among duplicated and not-duplicated mutations within large chromosomal gains.

One out of three patients whose sample was collected at relapse (IID_H198450) showed a clear presence of SBS25, a mutational signature previously linked to a still unknown chemotherapy agent. This signature has been observed in normal colorectal crypts in a patient previously treated with chemotherapy for non-Hodgkin lymphoma^33^ and in cHL cell lines^10^. This is the first report of SBS25 in relapsed cHL, demonstrating that similar to other cancers, cHL can acquire hundreds of mutations after exposure to distinct chemotherapy agents.^10,26,34-36^ The presence of a large SBS25 clonal mutational burden can be seen as a unique barcoding, reflecting the expansion of one single tumor cell that survived front-line treatment and subsequently took the clonal dominance.^35-37^ Leveraging this concept and estimating the SBS25 contribution before and after the three large chromosomal gains in this case, we observed chemotherapy-related mutations both before and after the large chromosomal gains. The presence of chemotherapy related mutations preceding large chromosomal gains indicates that these CNA events were acquired after exposure to chemotherapy.

### Molecular timing of multi-chromosomal gain events

cHL is known to be characterized by high ploidy and multiple chromosomal gains (e.g., WGD). Although most of these events are clonal, they may not have been acquired at the same time. To explore the temporal pattern of acquisition of chromosomal gains, including WGD, in cHL, we evaluated the corrected proportion of duplicated and non-duplicated clonal mutations within large chromosomal gains (i.e., molecular time approach).^23^ In line with our previous work,^24^ we restricted our analysis to clonal chromosomal trisomy, tetrasomy, gains, and copy neutral LOH larger than 1 Mb, with more than 50 clonal SNVs after having removed immunoglobulin loci and all localized hypermutated events (i.e., kataegis). Overall, in 62% gains the molecular time was higher than 0.5 (median 0.58; range 0.12-1.00), reflecting an intermediate-late acquisition pattern (**Figure 4a-b**).^23^ There were distinct events acquired particularly late, such as large gains on chromosome 2– *REL/XPO1*, 6q and 4q (**Figure 4c**). Other recurrent CNAs such as 9p24.1 – *PDCD1LG2/JAK2/CD274* were often intermediate. In 12/20 patients (60%) the final clonal copy number profile was acquired through at least two independent events (i.e., two groups of large gains with different molecular time, **Methods**; **Figure 4a-c** and **Supplemental Figure 6**). This suggests that HRS cells have a predisposition over time to acquire additional chromosomal gains.

**Figure 4.**
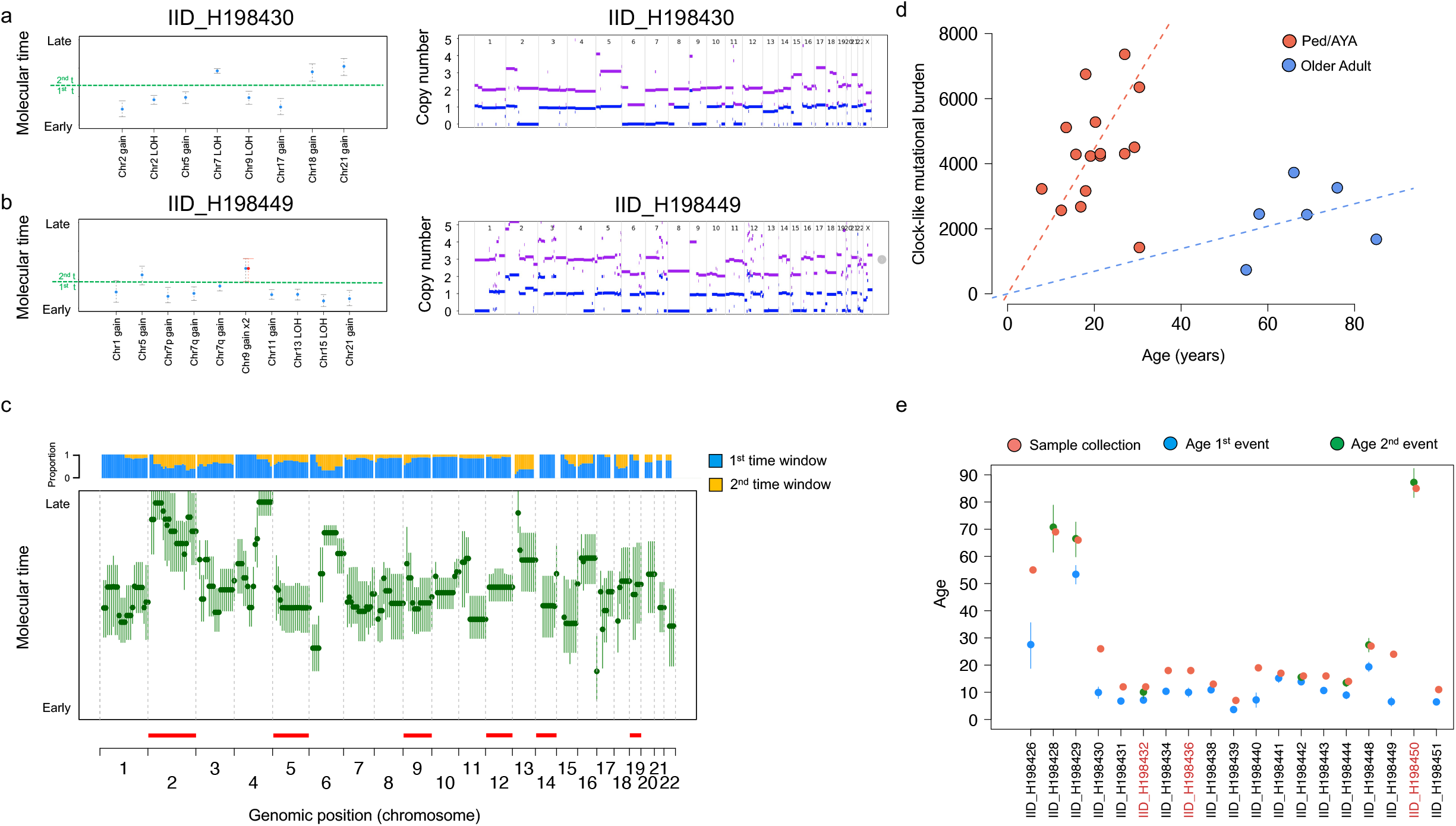
Molecular time and clock in cHL. **a-b)** Two examples showing molecular time estimates for two cases with multiple gains. Left, the molecular time (blue dots) estimated for each clonal gain and copy-neutral loss of heterozygosity with more than 50 clonal SNVs. Red dots represent the molecular time of a second gain occurring on a previous one. The dashed green line divides the two independent time windows in which different chromosomal gains were acquired (1^st^ t: first time window; 2st t: second time window. Right, standard copy number profile of 2 cHL cases. Horizontal purple and blue lines represent the total copy number and minor allele, respectively. **c)** Median molecular time estimates for chromosomal gains across each chromosome in cHL. Each chromosome was divided into 10 mb bins. Each green dot reflects the median molecular time across different patients with the gained bin. CI were generated using the median of CI molecular time estimate for each bin. Horizontal red line reflects GISTIC large amplifications. The blue/yellow plot represents the distribution of chromosomal gains acquired within either the first- or second-time window for each loci. **d)** Linear regression showing the association between age and SBS1/SBS5 mutational burden in cHL. **e)** Estimated patient age at the first (blue) and second (green) multi-gain event with 95% CIs. Red dots represent age at sample collection. Case numbers written in red represent cases collected at relapse.

To convert these relative estimations into absolute time (i.e., the age at which these events were acquired in each patient’s life), we utilized the clock-like mutational signatures as previously described.^10,25,26,30^ SBS1 and SBS5 (**Figure 3a-b**) have been reported to be universally present in all cells and to act in a constant rate over time.^10,23,30^ Based on this, we can quantify their activity before and after large chromosomal gains and convert the clock-like-based molecular time to an absolute value (i.e., years of life). We first confirmed that the SBS1 and SBS5 mutation rate were constant over time (**Figure 4d**). We observed a constant rate in pediatric and AYA patients as well as in older adults. Of note, the rates differed among age groups with a higher mutation rate in pediatric and AYA patients (p=0.01). Estimating the SBS1- and SBS5-based molecular time for large chromosomal gains acquired within the same time window and converting these relative estimates into absolute ones, we observed that the first multi-chromosomal gain event in cHL is often acquired several years before the diagnosis/sample collection (median lag between chromosomal gain and diagnosis: 5.3 years, range 1.8-27.4) (**Figure 4e**). In cases where we could assess the timing of the latest multi chromosomal gain events, it usually overlapped with the age at diagnosis/sample collection suggesting a potential role in the final lymphoma selection and clonal expansion.

### Structural variants and complex events

The high resolution of WGS allows us to perform the first characterization of the landscape of structural variants (SVs) in cHL.^38-40^ Applying *JaBbA* we were able to infer SV and CNA junction-balanced genome graphs with high fidelity allowing a detailed characterization of both single and complex events.^38^ Overall, we observed clear evidence of complex events such as chromothripsis (n=4), double minutes (*dm*, n=2), breakage-fusion-bridge (*bfb*; n=4), *pyrgo* (n=2), chromoplexy (n=1) and templated insertions (n=10) (**Figure 5a**). The role of single and complex events in the acquisition of CNAs involving distinct drivers emerged as heterogenous. For example, while most of the *PDCD1LG2/JAK2/CD274* high level gains were caused by large whole arm chromosomal gains or single events, in one patient these genes had multiple copies as consequence of *dm* (IID_H198450; **Figure 5b**). Similarly, in IID_H198427 a *bfb* event was responsible for the acquisition of 18 copies of *XPO1/REL* (**Figure 5c**).

**Figure 5.**
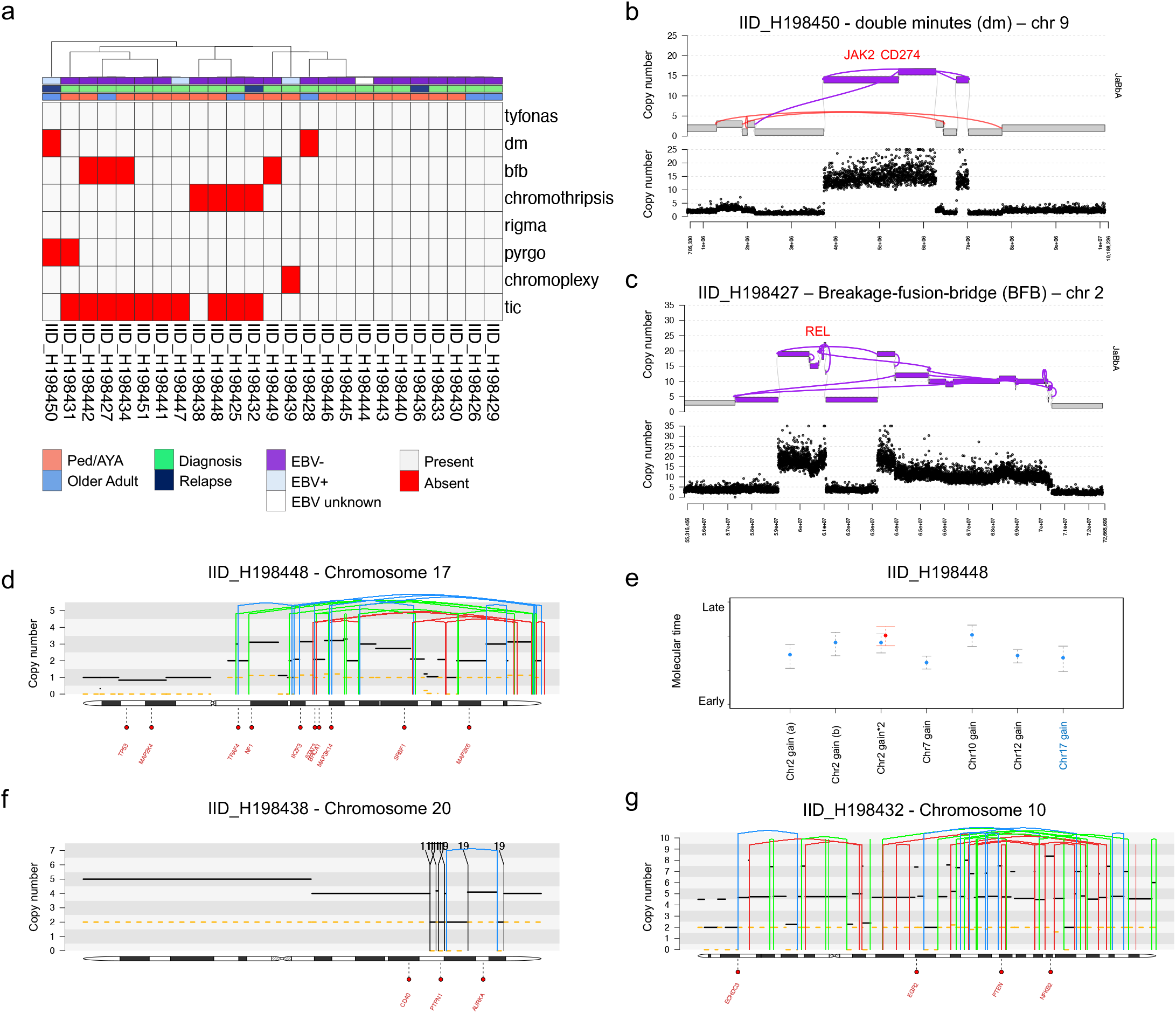
SV and complex event in cHL. **a)** Heatmap showing distribution of complex events across the 25 cHL cases evaluated by WGS. dm: double minutes, bfb: breakage-fusion-bridge, tic: templated insertion. **b-c)** Example of complex events involving key cHL drivers. **d-e)** Example of chromothripsis event in which it was possible to estimate the molecular time. Case IID_H198448 had an intrachromosomal chromothripsis event responsible for multiple large/intermediate chromosomal gain on chromosome 17 (**5d**). The molecular time of this event was similar to other gains acquired in the first/earliest-time window (**5e**). Based on this link we could estimate that that the chromothripsis event was acquired together with the other chromosomal gains (WGD) and approximately 10 years before the diagnosis (95% CI 8-13). **f-g)** Examples of two chromothripsis events that occurred before WGD. The two chromothripsis events were each responsible for a multiple copy number jump from 2:2 to 2:0. In **d), f)** and **g)**, the black and dashed yellow horizontal line represent the total number copy number and the minor allele, respectively. The blue, red, green, and black vertical lines represent inversion, deletion, tandem duplication, and translocation respectively. The partner of each translocation is reported on the top of the vertical black line.

To estimate the timing of loss-of-function events and the acquisition of distinct SVs, we utilized two approaches:^24^ 1) we linked SV breakpoints to the molecular time of chromosomal gains caused by the same SV (**Figure 5d-e**); 2) we estimated the relative time of SVs that occurred within large chromosomal gains based on the copy number of the SV breakpoint (**Figure 5f-g, Supplemental Figure 7**; see **Methods**). Applying these approaches, we were able to time the acquisition of single and complex events observing two relevant SV/CNA temporal patterns. In the first, chromothripsis emerged as an early event in cHL pathogenesis proceeding WGD in 3/3 patients in which this analysis was possible. (**Figure 5d-g**). In the second temporal pattern, we observed early acquisition of *PTPN1* deletion (i.e., before chromosomal gains/WGD) in 4/5 patients, suggesting the early driver role of this gene in cHL pathogenesis (**Supplemental Figure 7a** and **8**). Combining SV and SNVs in driver genes, *PTPN1* was involved in 36% (9/25) of cHL, either as early deletion (n=4) or SBS (n=5), each duplicated by a subsequent chromosomal gain.

### Using mutational signatures to identify the HRS cell-of-origin

The HRS cell-of-origin is suspected to be a B-cell that is unable to fully mature due to an unproductive B-cell receptor (BCR).^5,12,41-44^ This model has been supported by the detection of AID-mediated somatic hypermutation (SHM) on an unproductive VDJ, and nonsynonymous mutations involving AID off-target genes. To molecularly evaluate this model and the relationship between the GC and HRS cells, we explored the mutational signature within the immunoglobulin loci (Ig) and within the footprint of genes known to be involved by AID off-target activity (e.g., *PAX5, BCL6, XPO1*). These regions showed clear activity of AID/SBS84 (**Figure 3a** and **Figure 6a-c**). The presence of AID/SBS84 on Ig and on AID off-target genes has been reported in GC-derived malignancies.^10,26,45-47^ In these tumors, AID/SBS84 always co-occurs with a genome wide mutational process historically called “non-canonical” AID (SBS9) and thought to result from SHM. Contrary to this model, we did not observe any evidence of genome-wide SBS9 activity in AID/SBS84+ HRS cells (**Figure 6d**)^10,16,24,26,47-49^. This demonstrates for the first time that SHM is independent and temporally unrelated to SBS9.

**Figure 6.**
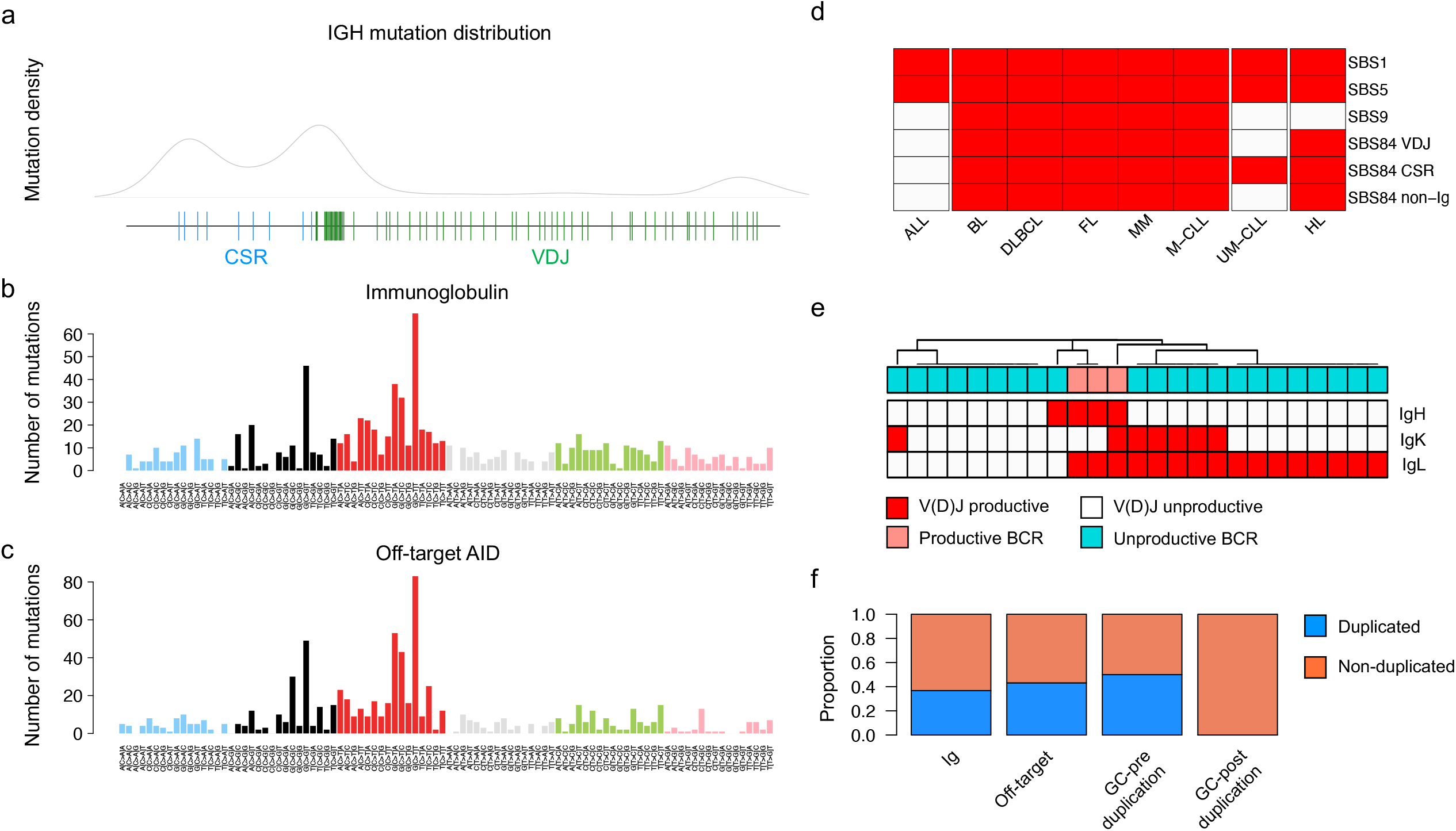
AID and SHM in cHL. **a)** Mutation density across the immunoglobulin loci. **b)** 96-mutational profile of all mutations occurred within the Ig loci. **c)** 96-mutational profile of all mutations occurred within known AID off-targets. The profile in **b)** and **c)** is identical to SBS84 (see Figure 3a). **d)** SBS9 and SBS84 distribution and activity across different lymphoproliferative disorders according to Maura et al.^17^ and Alexandrov et al.^10^). ALL: acute lymphoblastic leukemia; BL: Burkitt lymphoma; DLBCL: diffuse large B-cell lymphoma; FL: follicular lymphoma; MM: multiple myeloma; M-CLL: mutated IGHV chronic lymphocytic leukemia; U-CLL: unmutated IGHV chronic lymphocytic leukemia; HL: Hodgkin lymphoma. Each mutational signature activity is highlighted in red. **e)** Heatmap summarizing BCR productivity in our cohort of 25 cases with WGS. **f)** Proportion of duplicated and non-duplicated mutations within immunoglobulin loci and AID-off target involved by chromosomal duplications occurring within the earliest molecular time window. The last two bars on the right represent the expected proportion in the situation where the mutation was acquired before or after the chromosomal gain.

Next, to reconstruct the V(D)J of HRS cells, we ran *IgCaller*^50^ and confirmed that the VDJ rearrangements were unproductive in all but 3 patients (**Figure 6e, Supplemental Table 8**). To investigate when these unproductive BCRs were affected by SHM/AID, we focused on Ig and AID off-targets genes involved by large chromosomal gains. We observed both duplicated and non-duplicated AID mutations (**Figure 6f**) suggesting that, in the pre-cHL clone interaction with the GC precedes chromosomal duplications and, in some patients, might have been prolonged over time. Altogether this data molecularly confirms the cHL pathogenetic model in which a naive B-cell enters the GC, is exposed to AID and SHM, but as a consequence of an unproductive BCR fails to expand in the GC. The acquisition of distinct genomic drivers acquired before and during the GC reaction likely allows the pre-HL clone to escape negative selection in the GC (**Figure 7**). After being established and immortalized, in order to progress into cHL, the pre-cHL clone acquires additional driver events over time, in particular complex genomic events such as aneuploidies, WGD and SVs.

**Figure 7.**
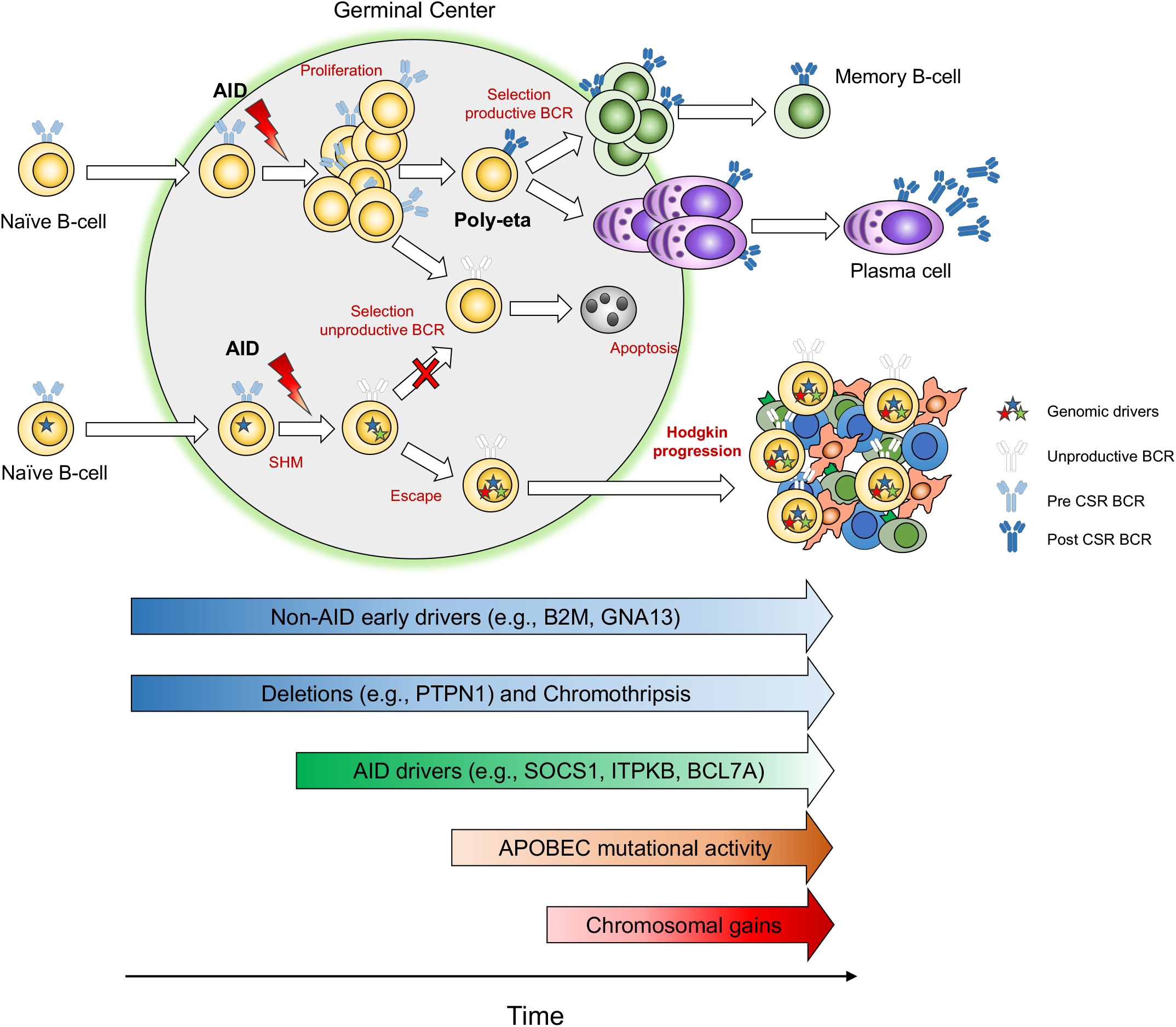
Summary of the proposed cHL germinal center-based pathogenetic model.

## Discussion

Human cancers are suspected to arise from a pre-malignant clone that evolves over time and often can be detected years before diagnosis, in some cases even in the peri-natal period.^23,25^ Understanding the chronology of mutational processes leading to malignancy can help guide novel diagnostic strategies and treatment approaches directed to early driver events. In this study, for the first time, we leveraged WGS resolution to elucidate a pathogenetic model for cHL whereby tumor development is shaped by the acquisition and selection of multiple drivers across different time windows over several years (**Figure 7**). In this model driver mutations in *B2M, BCL7A, PTPN1* and *GNA13* emerged as early events as did deletions in *PTPN1* and complex SV events such as chromothripsis. These early events largely occur before large chromosomal duplications which are often acquired at in intermediate/late timepoint which was determined to be still several years prior to diagnosis. This temporal pattern is different from what is observed in other hematological malignancies such as acute lymphoblastic leukemia, multiple myeloma and chronic lymphocytic leukemia, where large chromosomal duplications tend to be early events, potential playing a cancer-initiating role.^17,23,25,26,30,51^

We also observed several key differences in the mutational landscape of HRS cells across age groups. When compared to older adults age >40y, pediatric and AYA patients were found to have a higher mutational burden genome-wide, and an accelerated mutation rate in the clock-like signatures SBS1 and SBS5. This age segregation correlates with the two epidemiologic peaks of cHL which occur in AYAs age 15-40y and older adults age>55y, and suggests that the biology and the conditions in which cHL develop across these two peaks might be distinct. Additional studies are needed to investigate if the characteristics observed in younger patients with cHL may be due to accelerated B-cell aging.

The HRS cell mutational signature landscape revealed the first robust confirmation that the SBS9 signature represents a distinct GC mutational process independent from AID and SHM. The original term “non-canonical-AID” was first reported because, across the lymphoproliferative disorders tested (follicular lymphoma, CLL, multiple myeloma, Burkitt and diffuse large B-cell lymphomas), SBS9 always co-occurred with AID/SBS84 and SHM.^10,16,17,26,46,47^ This model has been recently challenged by the observations that: 1) AID/SBS84 can be active in CSR loci in CLL with unmutated immunoglobulin heavy-chain variable region gene in the absence of SBS9, 2) analysis of SBS9 genomic distribution in both tumor and normal B-cell genomes, and 3) by the fact that SBS9 profile seems to reflect polymerase-eta repair activity.^16,17,52,53^ In this study, the presence of all the GC hallmarks of SHM and AID activity with the absence of SBS9 in cHL fills an important gap in our understanding of which mutational processes are involved in both normal and pathological activity within the GC.

Lastly, with WGS comprehensive resolution, we confirmed and expanded the cHL pathogenic model proposed 30 years ago by Kupfer and collegues.^41-44^ Specifically, the interaction between pre-malignant cells and the GC emerged as a critical phase for cHL development. During this likely prolonged encounter, the pre-malignant cell survives and escapes the negative selection despite its unproductive BCR, potentially due to early and concomitant acquisition of genomic drivers. Overall, this study provides a critical new perspective on cHL pathogenesis and sheds light into which mutational processes are involved in the interaction between the GC and B-cells.

## Methods

### Sorting

The HRS, B and T cell sorting was performed essentially as describe previously.^2,7^ Briefly, single cell suspensions from CHL tumors containing up to 1 × 10^8^ cells were either taken fresh or rapidly defrosted at 37°C, washed in 50 mL of RPMI 1640/20% fetal bovine serum solution. The cells were stained with an antibody cocktail composed of CD64 FITC (22; Beckman Coulter (BC), Brea, CA), CD30-PE (HRS4; BC) CD5-BV510 (L17F12; Beckton-Dickinson (BD), San Jose, CA); CD40-PerCP-eFluor 710 (5C3; eBiosciences, San Diego, CA); CD20-PC7 (B9E9; BC); CD15-APC (HI98; BD); CD71 APC-A700 (YDJ1.2.2, BC); CD45 APC-H7 (2D1; BD), and CD95-Pacific Blue (DX2; Life Technologies, Grand Island, NY). All sorting experiments were performed on an FACS Aria-Fusion special-order research sorter using a 130-μm nozzle at 12 psi, acquiring up to 5 × 10^7^ cells and collecting HRS, B, and T cells from the tumor using a 3-way sort.

### Whole genome sequencing

Sample library construction was performed using the Kapa HyperPlus Kits with enzymatic fragmentation (Roche, Wilmington, MA). Fragmented gDNA were used to perform end repair, A-tailing and adapter ligation following the manufacturer’s instruction. The indexed library construct was split into two fractions, one fraction was used for WGS on NovaSeq6000 at PE2×150 cycles (Illumina, San Diego, CA) and another fraction was normalized and pooled at 4 samples from tumor samples and 4 samples for germline samples. Pooled sample libraries were hybridized with SeqCap EZ Human Exome v3.0 probes (Roche) for WES. The pooled, indexed and captured final libraries were used to sequence on Illumina HiSeq4000 sequencer at 2×100 cycles pair-end reads. The raw sequencing reads in BCL format were processed through bcl2fastq 2.19 (Illumina) for FASTQ conversion and demultiplexing for downstream data analysis.

### Processing of whole genome sequencing data

Overall, the median sequence coverage was 27.5X (range 15-55X; Supplemental Data 1). Short insert paired-end reads/FASTQ files were aligned to the reference human genome (GRCh37) using Burrows–Wheeler Aligner, BWA (v0.5.9). All samples were uniformly analyzed by the whole-genome analysis bioinformatic pipeline developed at the Memorial Sloan Kettering Cancer Center.^24,26,45^ Specifically: CaVEMan was used for SNVs, indels were analyzed with Pindel, CNAs were explored by Battenberg. To determine the tumor clonal architecture, and to model clusters of clonal and subclonal point mutations, the Dirichlet process (DP) was applied. BRASS and JaBba^38^ were used to detect SVs through discordantly mapping paired-end reads (large inversions and deletions, translocations, and internal tandem duplication). Complex events such as chromothripsis, chromoplexy, *dm, bfb*, templated insertions were defined and validated after manual inspection as previously described. ^39,40,54,55^ All SVs not part of a complex event were define as single. Immunoglobulin VDJ, HCDR3, CRS, and productivity were defined using *Igcaller*.^50^

### Whole-Exome Sequencing

Whole exome sequence data reads were aligned to the reference human genome (GRCh37) using the Burrows–Wheeler Alignment tool (bwa mem v0.7.12). PCR duplicate read removal, InDel realignment, fixing mates and base quality score recalibration was applied to the aligned bams using PICARD tools or the Genome Analysis Toolkit (GATK) according to GATK best practices. Samples in the selected cohort had an average coverage of 76X in tumors and 49X in germline (range 15X - 135X). The tumor purities were inferred using TITAN and ranged between 31-91%. WXS somatic calls were performed using CaVEMan for SNVs, Pindel for indels, and Facets for CNA.

### Mutational signatures

Analysis of SBS signatures was performed following our published workflow based on three main steps:^17^ 1) de novo extraction; 2) assignment; and 3) fitting.^17^ For the de novo extraction of mutational signatures we ran two independent algorithms; *SigProfiler* and the hierarchical Dirichlet process (hdp).^10,26^ Next, each extracted process was assigned to one or more mutational signatures included in the latest COSMIC v3.2 catalog (https://cancer.sanger.ac.uk/signatures/sbs/). Lastly, *mmsig*, a fitting algorithm designed for hematological cancers (DOI: 10.5281/zenodo.4541703), was applied to accurately estimate the contribution of each mutational signature in each sample. ^29^

### Timing copy number and structural variant events

The relative timing of each multi-chromosomal duplication event was estimated using the R package *mol_time* (DOI: 10.5281/zenodo.4542145).^24,26^ This approach allows the estimation of the relative timing of acquisition of large (>1Mb) and clonal chromosomal gains (3;1 or 4:1), trisomies (3:1), copy neutral LOH (2:0), and tetrasomies (4:1) using the purity-corrected ratio between duplicated clonal clonal mutations (defined by DP). SNVs were defined as duplicated or not according to the VAF corrected for the cancer purity which was estimated combining purity estimates from both Battenberg and from the SNV VAF density and distribution within clonal diploid regions. Overall, after removing immunoglobulin loci and kataegis events, only CNA segments with more than 50 clonal mutations were considered.^24,26^ Tetrasomies, with both alleles duplicated (2:2), were removed given the impossibility of defining whether the two chromosomal gains occurred in close temporal succession or not.^24,26^ Using this approach, we were able to estimate the relative molecular time of each gained CNA segment allowing to define if different chromosomal gains were acquired in the same or different time windows. To define if different gains occurred in one single time window or in different independent events we used multiple hierarchical clustering approach for each single bootstrap solution (*hclust* R function; *www.r-project.org*) and we integrated the most likely results with the Battenberg CNA changes over the time.

Next, we convert the relative molecular time estimate into an absolute estimate (i.e., years of life) estimating the molecular time using only the SBS1 and SBS5 mutational burden within large chromosomal gains acquired in the same time window. Applying this workflow we converted the SBS1 and SBS5-based molecular clock into an absolute time for the acquisition of these events in each patient’s life.^23,25,26^ Confidence of intervals were generated by bootstrapping 1000 times the molecular time estimate. Similar to the relative molecular time approach, only multi-gain events with more than 50 SBS1 and SBS5 clonal mutations were included.

To estimate the timing of loss-of-function events and the acquisition of distinct SVs, we utilized two previously reported approaches:^24^ 1) we linked SV breakpoints to the molecular time of chromosomal gains caused by the same SV; 2) we estimated the relative time of SVs that occurred within large chromosomal gains based on the copy number of the SV breakpoint. Approach (2) is based on the concept that any SV involving a large gain can produce three different copy number scenarios: i) SV associated loss occurred on the allele involved by the duplication before the duplication. In this scenario, the deleted segment will not be duplicated creating a copy number jump of at least 2 copies with the adjacent duplicated segments (e.g., 3:1 and 1:0).; ii) the SV is responsible for a loss (or gain) on one of the two duplicated alleles (e.g., from 2:1 to 1:1), suggesting that the SV event occurred after the chromosomal gain. iii) SV causes the loss (or gain) of part of the minor allele creating a copy neutral LOH (e.g., from 2:1 to 2:0;. In this last scenario it is not possible to establish if the SV occurred before or after the chromosomal duplication.

The recurrent mutations in driver genes chronological acquisition order was estimated combining the Battenberg cancer cell fraction and pre/post gain data into a Bradley Terry model as previously described, including just the earliest sample of each patient ^24^

### Combined Immunophenotyping and FISH analysis

After immunophenotyping with mouse anti-CD30 antibody labelled with AMCA (blue), the relevant tissue slides with coverslip are briefly reviewed and recorded under fluorescence microscope for the intensity and quality of CD30 staining. After removing the coverslip, the slides are washed in 2xSSC at room temperature for 5 minutes and then fixed with 10% neutral buffered formalin for 20 minutes, followed by dehydrating five minutes each in a series of 70%, 85% and 100% of alcohol. Slides then go through the pretreatment for 10 minutes with 20 mM Citrate Buffer/1% NP-40 Mixture, pH 6.0-6.5, followed by protease treatment at 40º C for 10-15 minutes (Abbott Molecular, Des Plaines, IL), then by dehydrating 2 minutes each in a serios of 70%, 85% and 100% of alcohol. Relevant FISH probes were selected based on WGS ploidy findings and primarily were those used in clinical diagnosis tests, including CDK2 and CEP12, PDL1 and PDL2 and CEP9, and 19p and 19q, and EWSR1 (Abbott Molecular, Des Plaines, IL and Genomic Empire, Buffalo, NY). These probes were labeled in spectrum orange, green or aqua, respectively. After applying the FISH probes to the tissue areas, both tissue and probes were co-denatured at 94º C for 7 minutes, and then incubated at 37º C overnight, followed by post-hybridization washing in 2xSSC/0.3% NP-40 at 77º C for one minute. Tissue sections were counterstained with antifade medium without DAPI (Vector Laboratories, Burlingame, CA).

The slides are evaluated under fluorescence microscope coupled with appropriate filters for CD30 immunophenotype and for the relevant probes. Signal analysis was performed in combination with tissue structure and cell morphology correlation and was focused on the interested tissue areas with strong CD30 positive cells only. The copy numbers of individual probes were counted in each case.

### Data analysis and statistics

Data analysis was carried out in R version 3.6.1. Standard statistical tests are mentioned consecutively in the manuscript while more complex analyses are described above. Wilcoxon rank-sum test between three groups was run using *pairwise*.*wilcox*.*test* R function with all p value adjusted for FDR. All reported p-values are two-sided, with a significance threshold of < 0.05.

## Acknowledgements

This work was supported by the Children’s Oncology Group, the Hartwell Foundation, the Gant Family Foundation, the Sylvester Comprehensive Cancer Center NCI Core Grant (P30 CA 240139), and the Memorial Sloan Kettering Cancer Center NCI Core Grant (P30 CA 008748). F.M. is supported by the American Society of Hematology. F.M. and O.L. are supported by the Riney Family Foundation. L.G.R is supported by the NIH (K08CA219473) and the Hartwell Foundation.

## Author contributions

F.M., M.R., L.G.R. designed and supervised the study, collected and analyzed the data and wrote the paper. M.I. J.Z.X, B.B, B.Z., F.A., K.H.M., K.W.E., M.U., F.H, W.Z., Q.G., V.Y., O.L., C.M., E.C., T.H., O.E. collected and analyzed data. S.P., M.O., E.R., M.L., G.W., C.F. and M.B. provided patient samples. All authors reviewed and edited the manuscript.

## Data availability

The other already published data are deposited in the EGA and dbGap database under the following accession numbers: EGAS00001001692: PCAWG cohort [https://ega-archive.org/studies/EGAS00001001692]; EGAD00001003309, 67 WGS raw data from 30 patients with MM and myeloma precursor conditions [https://www.ebi.ac.uk/ega/datasets/EGAD00001003309]; phs000348.v2.p1, 22 WGS raw data from patients with MM [https://www.ncbi.nlm.nih.gov/projects/gap/cgi-bin/study.cgiãstudy_id=phs000348.v2.p1]. All these data are available under restricted access, access can be obtained by contacting the public depository.

