## Supplemental Figures for "Molecular evolution of classic Hodgkin lymphoma revealed through whole genome sequencing of Hodgkin and Reed Sternberg cells"

### The genomic history of classic Hodgkin lymphoma

#### Supplemental Figure

**Supplemental Figure 1. Amplification-based mutational artifacts**. The 96-mutational profile of the identified palindromic artifact. b) IGV screenshot of one palindromic artefactual mutation.

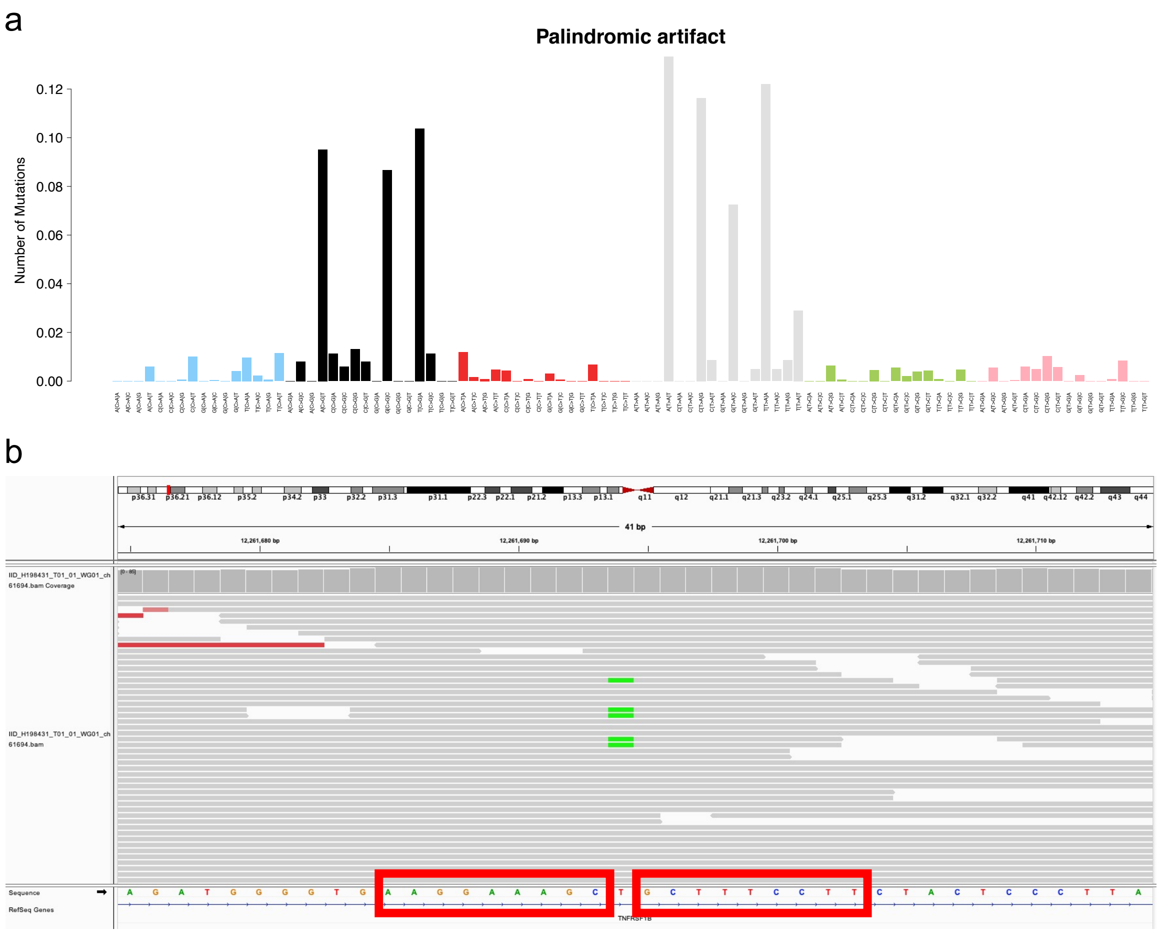

**Supplemental Figure 2. AID/SHM is responsible for *SOCS1*, *ITKB* and *BCL7A* multiple mutations in the same patients.** A completely different 96-mutational profile was observed for *B2M* and *GNA13*, suggesting that other mechanisms are responsible for multiple hits on this gene.

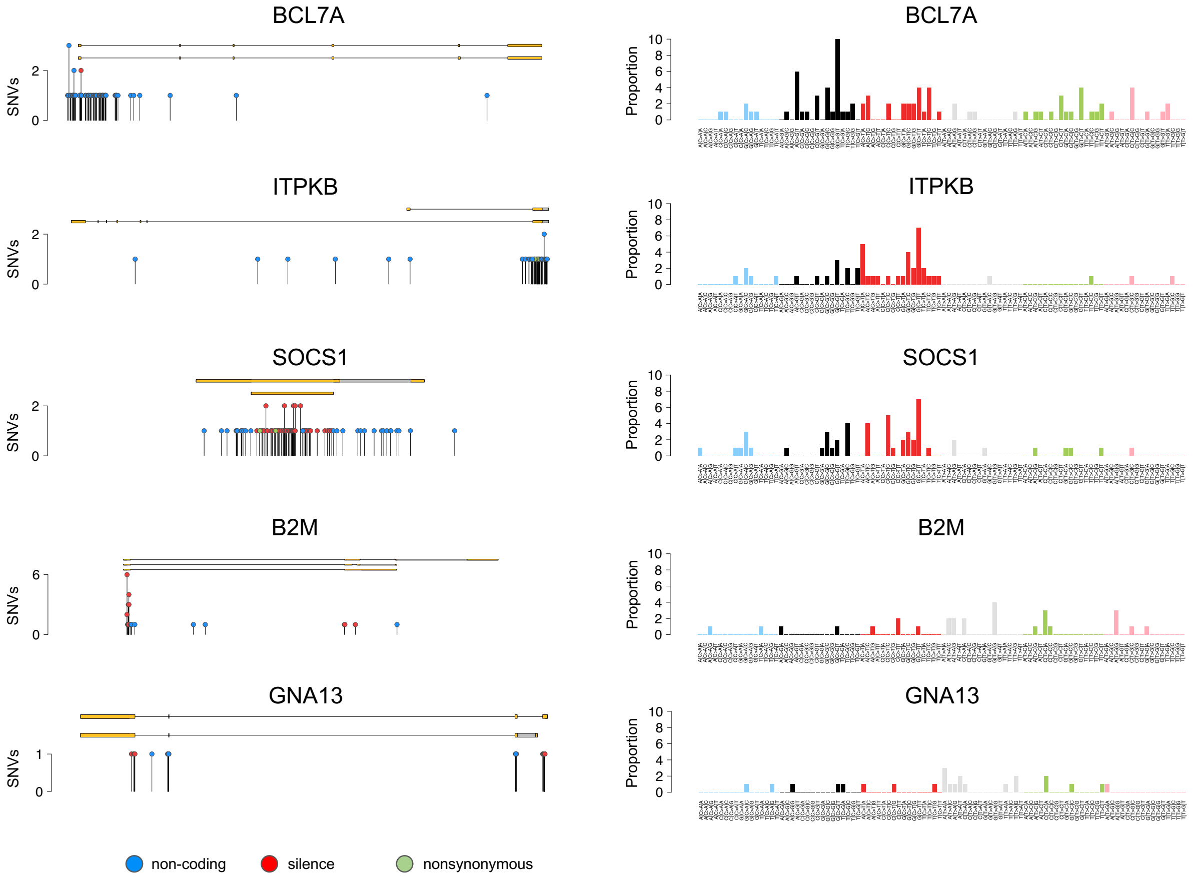

**Supplemental Figure 3. Combined CD30 immunophenotyping and FISH analysis determine the ploidy of HRS cells in each case.** FISH analysis of the relevant chromosomal targets, i.e., *CDK4* (12q14) and *CEP12* (centromere probe), *PDL1* and *PDL2* (9p24.1) and *CEP9*, 19p13 and 19q13, and *EWSR1* (22q12) show the copy numbers of at least three chromosomes and determine if the HRS cells are diploidy with an average of two signals for each target (IID_H198425) or triploid/tetraploid with three to four signals of the relevant probes (IID_H198426). CD30 positive cells are stained blue, and FISH probes in each test are labeled spectrum orange, spectrum green or spectrum aqua, respectively.

IID_H198425 – no whole genome doubling

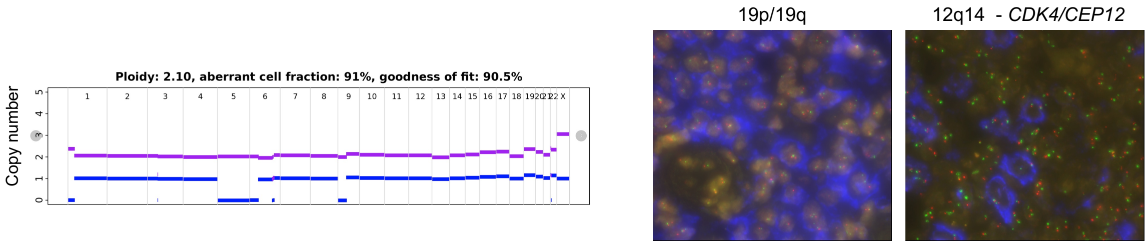


IID_H198426 – whole genome doubling

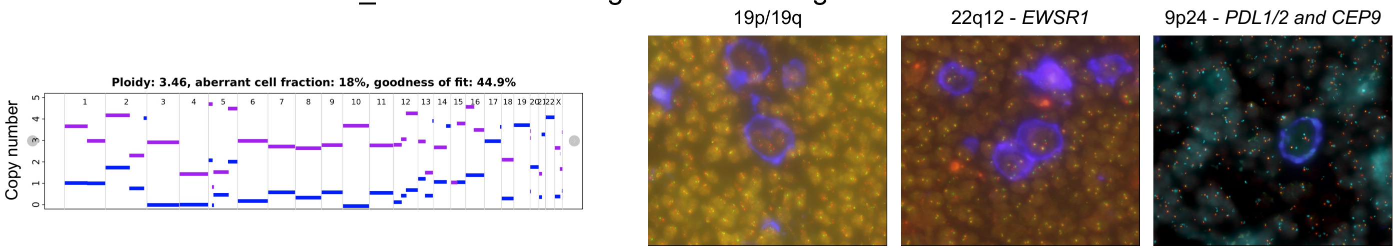


n

**Supplemental Figure 4. Cumulative copy number plot for each significant driver genes identified by GISTIC2.0.** The red vertical line represents the gene start and end, the grey dashed lines represent the start, end and centromere of the plotted chromosome.

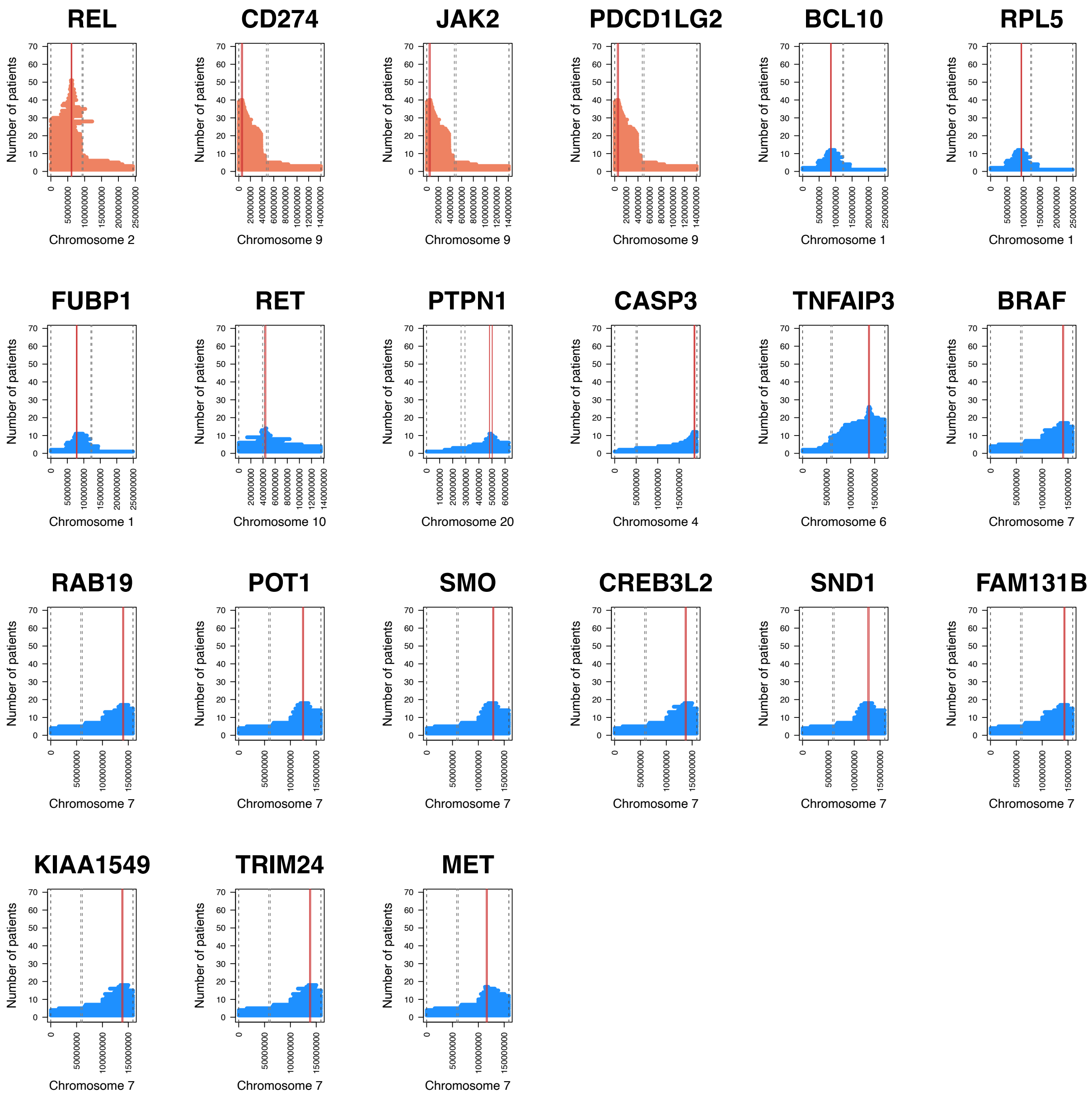

**Supplemental Figure 5. Size of copy number alteration (CNA) involving gene drivers within GISTIC peaks.** The green dashed line represents 10 Mb threshold used to distinguish large and focal CNA events.

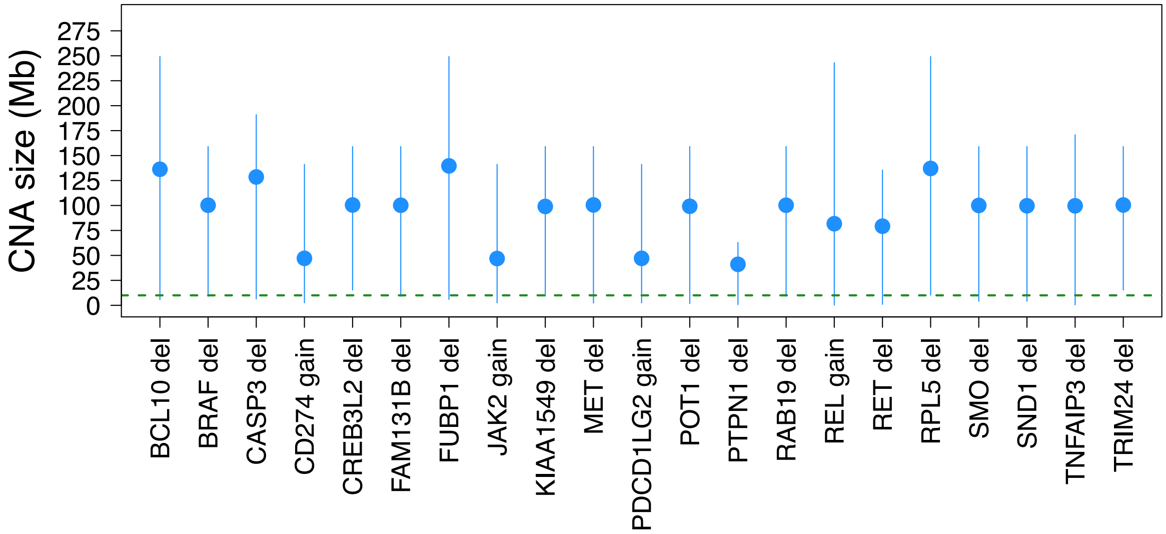

**Supplemental Figure 6. Molecular time estimates for 22 cases with multiple chromosomal gains.** Blue dots represent the molecular time estimate each clonal gain and copy-neutral loss of heterozygosity with more than 50 clonal SNVs. Red dots represent the molecular time of an eventual second extra gain occurred on a previous one. Horizontal dashed green line separates independent time windows in which chromosomal gains were acquired.

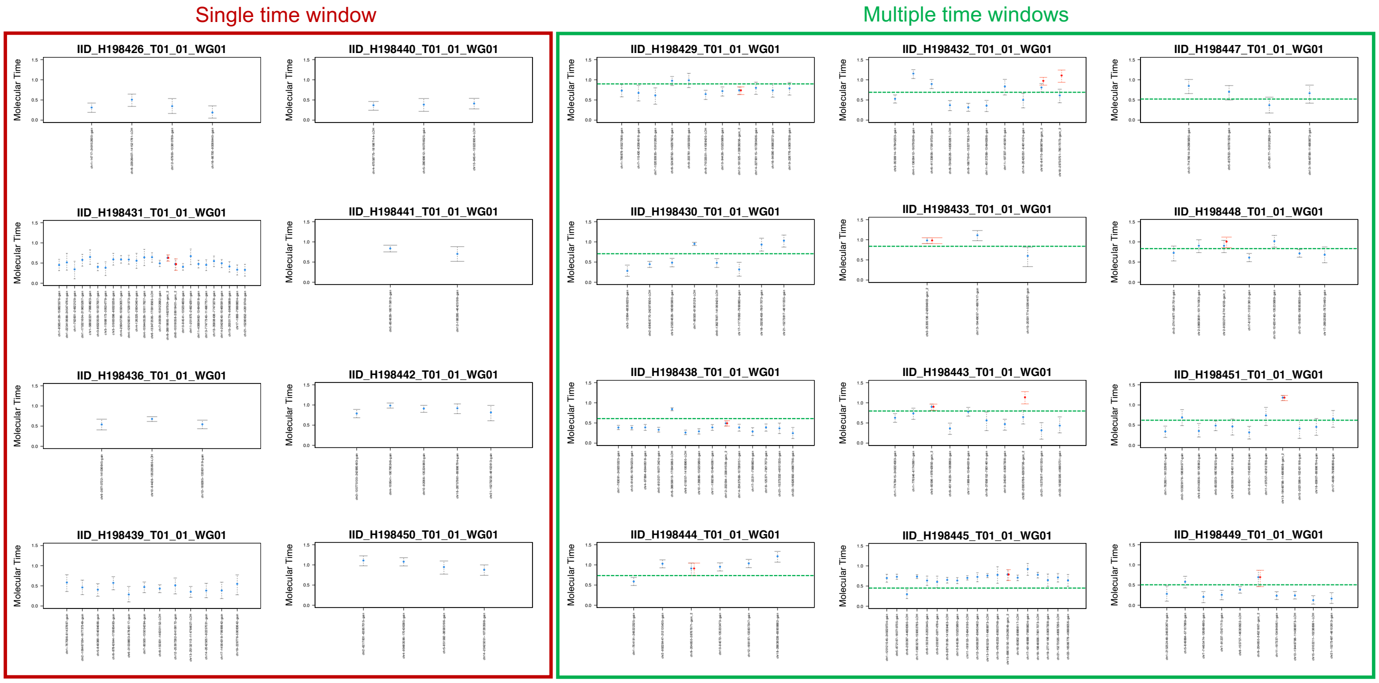

**Supplemental Figure 7. Three examples and schemes that summarize our approach to time SV within large chromosomal gains.** In a), c) and e) the black and dashed yellow horizontal line represent the total copy number and the minor allele, respectively. The blue, red, green, and black vertical lines represent inversion, deletion, tandem duplication, and translocation respectively. The partner of each translocation is reported on the top of the vertical black line. In b), d) and f) the scheme of the copy number timing for SV events showed in a), c) and e), respectively.

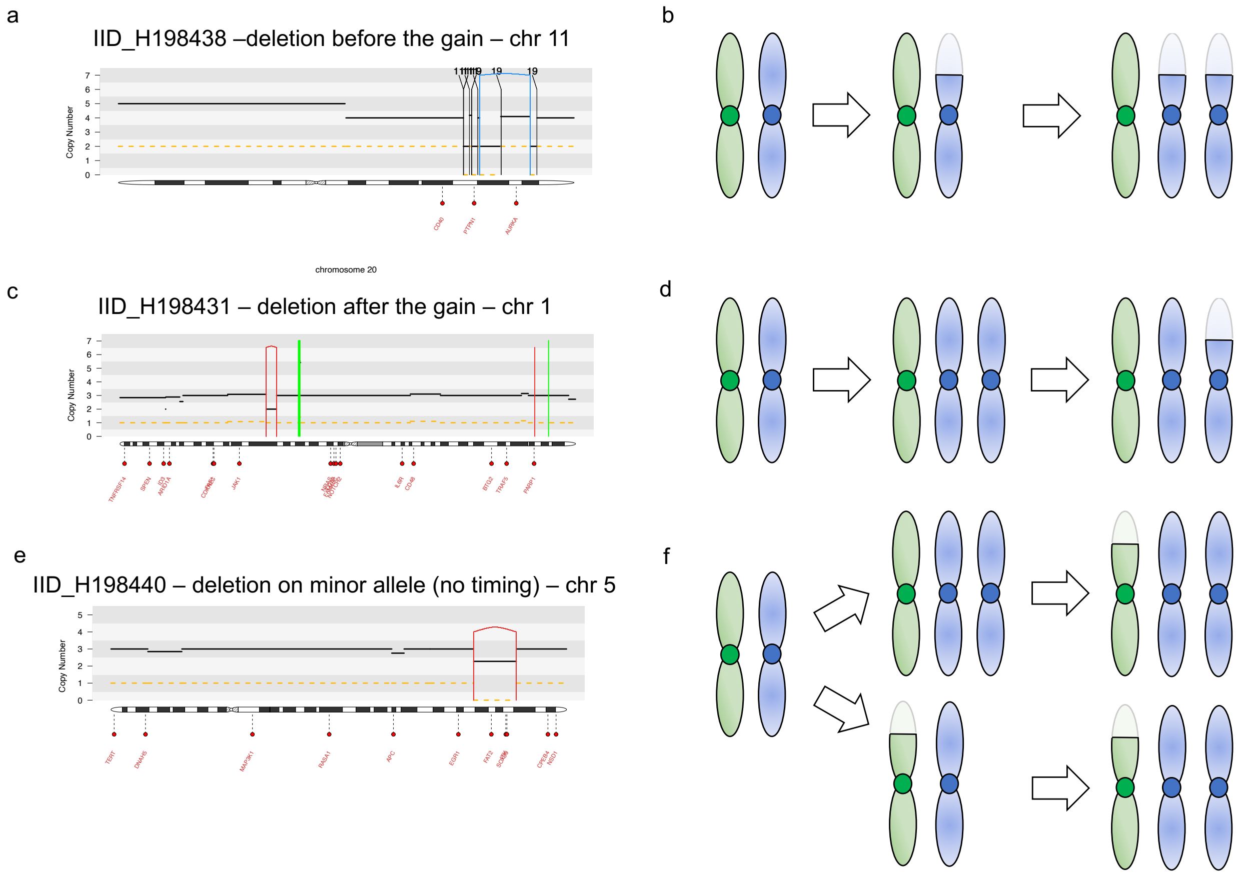

**Supplemental Figure 8. Timing of *PTPN1* deletion in cHL.** In each plot the black and dashed yellow horizontal line represent the total copy number and the minor allele, respectively. The blue, red, green, and black vertical lines represent inversion, deletion, tandem duplication, and translocation respectively. The partner of each translocation is reported on the top of the vertical black line.

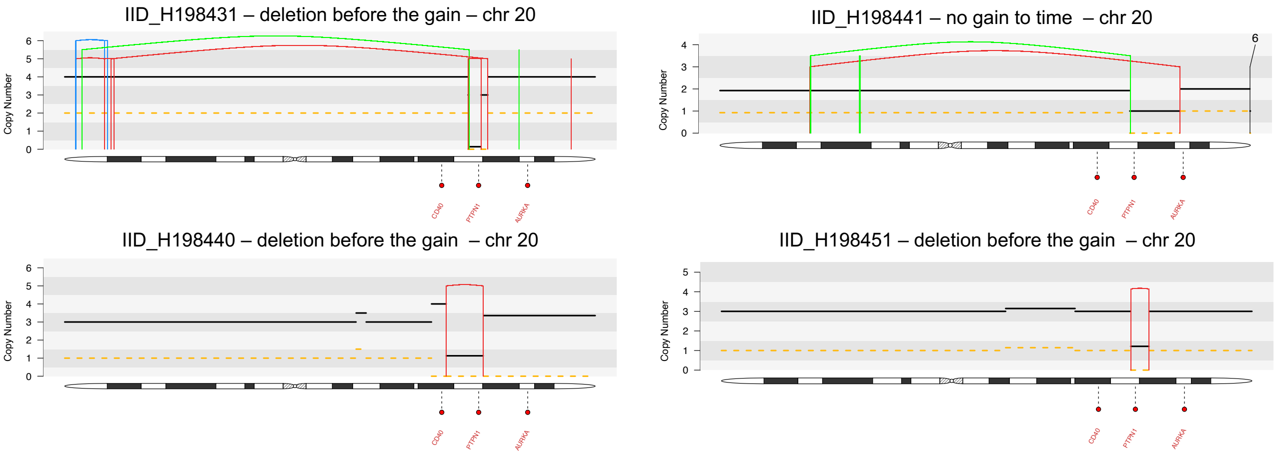
